# End-to-end Bayesian analysis of ^14^C dates reveals new insights into lowland Maya demography

**DOI:** 10.1101/2020.07.02.185256

**Authors:** Michael Holton Price, José M. Capriles, Julie A. Hoggarth, Kyle Bocinsky, Claire E. Ebert, James Holland Jones

## Abstract

Archaeologists and demographers increasingly employ aggregations of published radiocarbon (^14^C) dates as demographic proxies summarizing changes in human activity in past societies. Presently, summed probability densities (SPDs) of calibrated radiocarbon dates are the dominant method of using ^14^C dates to reconstruct demographic trends. Unfortunately, SPDs are incapable of converging on their true generating distributions even as the number of observations gets large. To overcome this problem, we propose a more principled alternative that combines finite mixture models and Bayesian inference to identify the generating distribution of a set of radiocarbon dates. Numerical simulations and an assessment of the statistical identifiability of our method demonstrate that it correctly converges on the generating distribution. We apply this novel end-to-end Bayesian approach to reconstruct prehistoric Maya demographic growth using a recently compiled Mesoamerican radiocarbon database. Our results show that the Maya Lowlands experienced a century of rapid growth rates (1%) during the Late Classic, followed by a rapid decrease in population during the Terminal Classic, and a subsequent more-modest resurgence in population during the Postclassic. Additionally, a detailed population reconstruction of the important political center of Tikal verifies that slow population growth between the Preclassic and Early Classic gave pace to rapid growth starting around AD 500 and peaking at the beginning of the eight century. Our proposed method verifies previous reconstructions based on settlement patterns and ceramics, but with far more precise time-resolution and characterization of uncertainty than has been possible.

## Introduction

Since John Rick proposed that sets of published radiocarbon (^14^C) dates from a region provide a useful proxy for relative population size through time^1^, an increasing number of studies have been published using ^14^C data sets from across the world to track and reconstruct human demographic change. Indeed, over two hundred articles have been published on the topic since 2010, showing the still-growing impact and influence of this approach for studying past population dynamics^2–4^. Despite the popularity of using radiocarbon “dates as data,” important criticisms to the approach have emerged (and many have been addressed)^5–10^. Broadly, these criticisms can be divided into two categories: (a) the radiocarbon dates available for study are not representative of past population sizes, and (b) even if they are representative, the interaction of the radiocarbon calibration curve with measurement uncertainty hampers inference and the conclusions one can make. We will refer to the former as the “bias problem” and the latter as the “summary problem”^11^.

The current, dominant method to summarizing sets of radiocarbon dates is to construct summed probability distributions or densities (abbreviated as SPDs). To generate SPDs, each individual observation is first calibrated, which involves adopting a prior probability for the calendar date of each observation usually a uniform prior – and updating this prior to yield a posterior density from the likelihood of obtaining the radiocarbon determination, given the known radiocarbon calibration curve and measurement uncertainty. Next, these individual posterior densities are summed, and the resulting summed curve is normalized by dividing by the number of observations, yielding a density distribution that integrates to one. The resulting SPD is then typically used as either a rough equivalent of a demographic growth curve, compared to specific growth models to determine statistically significant deviations, and/or compared to various sequences of paleoclimatic and paleoenvironmental change^12–14^.

Crucially, because the calibration is done independently for each observation, the SPD incorporates measurement error and ambiguity arising from the radiocarbon calibration curve only at the level of the individual sample. Hence, SPDs, and all approaches based on them, do not fully use the information available in a set of radiocarbon dates. It is straightforward to show that, even as the number of observations gets large, the SPD does not converge on the distribution that generated a set of radiocarbon samples (Figure 1). The alternative use of kernel density estimation plots (KDEs) produces results less affected by the high frequency noise of the calibration plots but is still largely unable to deal with the dispersion of the data produced by the measurement and calibration uncertainty^11^. In the next section, we describe an end-to-end Bayesian algorithm that does not have these failings and, furthermore, supports statistical inference and hypothesis testing that is difficult or impossible with SPDs or KDEs.

**Figure 1.**
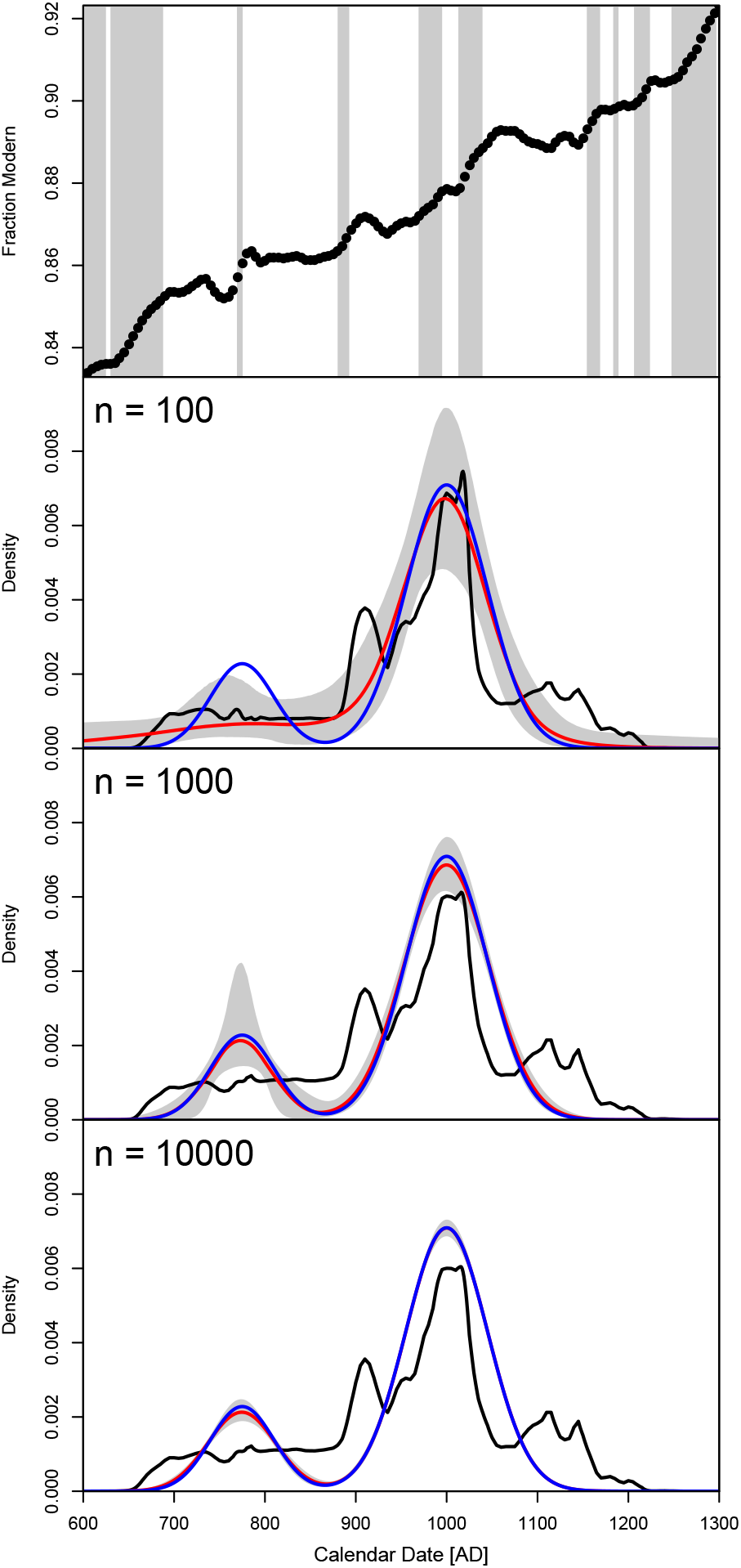
Summary of simulation results. The top figure shows the radiocarbon calibration curve fraction modern as a function of calendar date; grey bands show where the fraction modern has a unique corresponding calendar date. The next three figures show in turn the simulation results for N=100, N=1000, and N=10,000. The blue curve is the true, target density (identical across simulations). For each simulation, the red curve shows the 50% quantile from the end-to-end Bayesian sampling, and the grey bands delineate the 2.5% and 97.5% quantiles; the black curve shows the SPD. As *N* increases, the end-to-end Bayesian approach correctly reconstructs the target distribution, whereas the SPD converges on a stable shape that does not reconstruct the target distribution.

## End-to-End Bayesian Modeling

In this article, we advocate an end-to-end approach in which Bayesian inference is done on alternative demographic models. More precisely, we assume that the set of radiocarbon dates to be summarized is drawn from a single, parameterized probability distribution, and the likelihood function used to update the Bayesian priors is defined on the entire set of radiocarbon dates simultaneously. Doing such end-to-end inference more effectively accommodates the equifinality induced by the radiocarbon calibration curve arising from fluctuating atmospheric ratios of ^14^C to ^12^C. In contrast, summed probabilities and all approaches based on them accommodate the ambiguity caused by equifinality independently for each sample prior to “fusing” data via a sum. From an information theoretic standpoint, any inferential method that starts with summed probabilities eliminates potentially useful information. The reason is that addition (summation) is a logically irreversible process: 2 + 2 and 1 + 3 both equal 4, so if we only know the result of this sum there is no way we can reconstruct the original inputs. Similarly, summing the posterior densities of individual radiocarbon samples is an irreversible step, and measurement uncertainty combined with equifinality in the radiocarbon calibration curve can cause two distinct sets of dates to have similar summed probability curves.

An unfortunate aspect of SPDs, which arises in part from the intrinsic irreversibility of summation, is that even with an infinite number of observations, SPDs cannot reproduce the original distribution that generated the radiocarbon dates. In contrast, the end-to-end Bayesian approach we introduce in this article can. We validate our approach using two lines of evidence. First, we utilize simulation studies involving finite Gaussian mixtures to show that with enough samples, the generating distribution is exactly reconstructed. Second, we introduce a linear-algebra based method for analyzing the statistical identifiability of parametric models of the radiocarbon measurement process. Briefly, a parametric model is identifiable if two distinct values of the parameter vector necessarily yield different probability densities for the measurements one can make. We show that finite Gaussian mixtures are identifiable even with the intervening effects of measurement error and calibration uncertainty.

We now describe in detail our end-to-end Bayesian algorithm, including the specification of prior probabilities and the likelihood calculation. The algorithm is available in an open source R package, **baydem**, available at https://github.com/eehh-stanford/baydem. The simulations and analyses in this article can be generated using the code at https://github.com/eehh-stanford/price2020, which also provides a template for using the **baydem** package.

Let *t* be calendar date and let *θ* be a vector that parameterizes the density function *p*(*t*|***θ***) from which a set of (unobserved) calendar dates are sampled; collectively, we represent the data available to infer these dates by *D*. The posterior probability of *θ* is

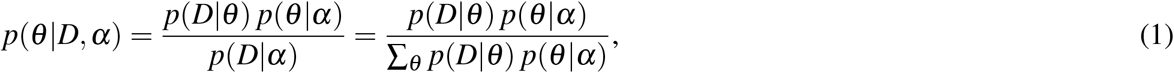

where the vector *p*(*θ* |*α*) parameterizes the prior probability.

### Parameterization of density

Finite Gaussian mixtures provide a highly flexible class of density around which to build our model^15^. We parameterize the density as a mixture of Gaussians with *K* components,

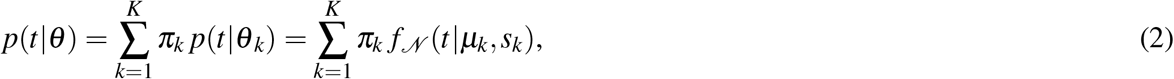

where 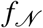 is the Gaussian probability density, *π*_*k*_ the mixture proportion, *μ*_*k*_ the mean, and *s*_*k*_ the standard deviation. For Bayesian inference, the density is normalized to integrate to 1 on the truncated interval *τ*_min_ to *τ*_max_.

### Parameterization of prior

The full parameterization vector of the Gaussian mixture described in the preceding section is

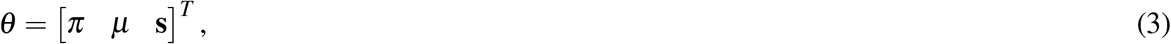

where [*π* = *π*_1_ … *π*_*K*_]^*T*^ is the column vector of mixture components (and similarly for *μ* and **s**). We assume that *π* is drawn from the symmetric Dirichlet distribution, *π* ~ dirichlet(*α*_*d*_, *K*), where *α*_*d*_ is a scalar concentration parameter, and that the means are drawn uniformly on the interval *t*_min_ to *t*_max_, *μ*_*k*_ ~ uniform(*τ*_min_, *τ*_max_). We assume the standard deviations are drawn from a gamma distribution, *s*_*k*_ ~ gamma(*α*_*s*_, *α*_*r*_), where *α*_*s*_ is the shape parameter and *α*_*r*_ is the rate parameter. For the simulation, we use *α*_*d*_ = 1, *α*_*s*_ = 3, *α*_*r*_ = (*α*_*s*_ − 1)*/*300 = 0.00667 (a maximum at 300 years), *τ*_min_ = 600, and *τ*_max_ = 1300 (dates are AD). For the Maya data, which span a greater range of calendar dates than the simulated data, we use *α*_*d*_ = 1, *α*_*s*_ = 10, *α*_*r*_ = (*α*_*s*_ − 1)*/*500 = 0.018 (a maximum at 500 years), *τ*_min_ = −1100, and *τ*_max_ = 1900.

### Likelihood for radiocarbon measurements

Let *ϕ*_*m,i*_ be the fraction modern (ratio of carbon-14 to carbon-12 normalized such that a sample from 1950 yields a value of 1) for measurement *i* and *σ*_*m,i*_ the corresponding uncertainty. The uncalibrated calendar date and associated uncertainty are, respectively,

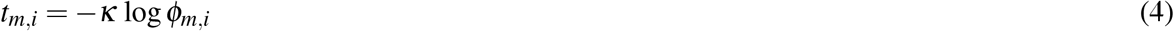

and

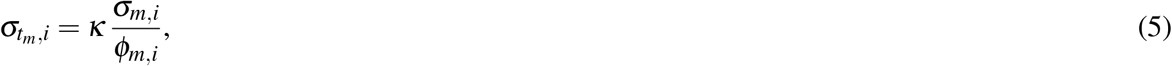

where *κ* = 8033 is the reference decay rate of carbon-14. The probability of the measurement *i* given the calendar date *t* is

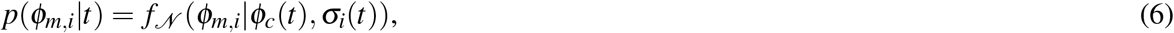

where

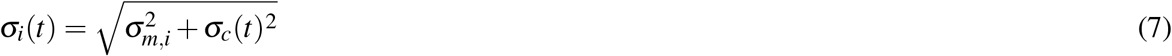

is the total uncertainty, accounting for both measurement and calibration error, *ϕ*_*c*_(*t*) is the calibration curve value (fraction modern), and *σ*_*c*_(*t*) is the associated calibration uncertainty. The likelihood of measurement *i* is

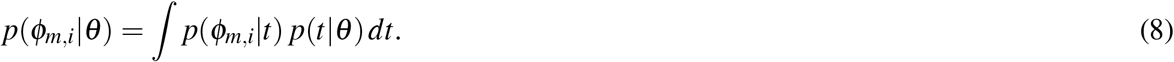

Equation 8 elides the dependence on the calibration curve and measurement uncertainty. We approximate this integral with a Riemann sum at the grid points

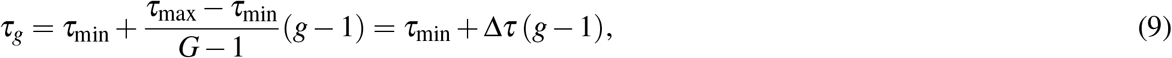

where *g* = 1, …, *G* indexes grid points and Δ*τ* is the grid spacing. The Riemann sum is

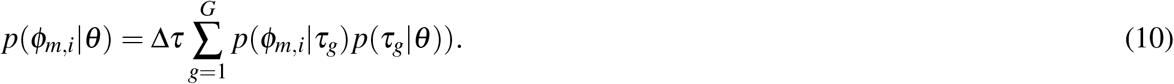

Define the measurement matrix as (using index notation)

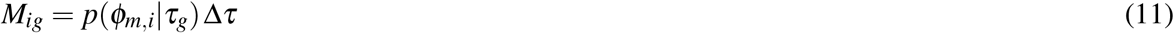

and define the sampling vector as

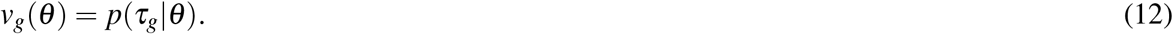

The vector of likelihoods *h*_*i*_ is

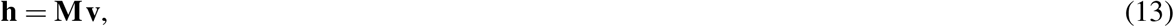

where **h** and **v** (but not **M**) depend implicitly on *θ*. The overall likelihood is the product of elements of **h**

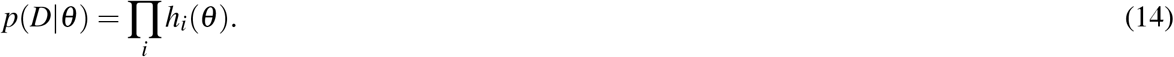

### Sampling the posterior

Since the posterior density cannot be directly calculated, we sample from the posterior using the Hamiltonian MCMC algorithm implemented in Stan. For some observable that depends on calendar date, *ψ*(*t*|*θ*), the posterior estimate is

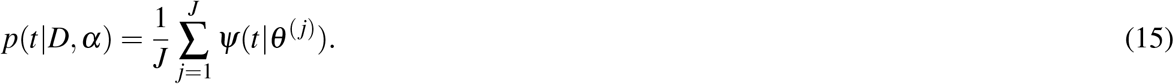

where *θ*^(*j*)^ is the *j*-th MCMC sample. The set of samples *ψ*(*t*|*θ*^(*j*)^) can be ordered to calculate quantiles. For density (e.g., Figure 1) *ψ*(*t*|*θ*) = *p*(*t*|*θ*) and for rate (e.g., Figure 4) 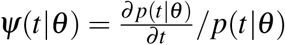.

## Model Validation

### Simulations

To demonstrate the potential of the method and assess the accuracy of the end-to-end Bayesian inference, we run a series of simulations for 100, 1000, and 10,000 radiocarbon samples and compared them to SPDs (Figure 1). The true, target distribution (blue curve) is a two-component Gaussian mixture for which the mixture proportion, mean, and standard deviation of each component are, respectively, 0.2 / AD 775 / 35 years and 0.8 / AD 1000 / 45 years. We truncated the distribution to be on the interval AD 600 to AD 1300 (which had a vanishing effect since the two component Gaussians have a vanishing probability density at AD 600 and AD 1300). For each simulated observation, we first sampled randomly for the calendar date from the target density function. Second, we simulated the data-generation process by: (a) calculating the mean fraction modern using the radiocarbon calibration curve (IntCal13)^16^ and then (b) making a random draw for the error of the measurement. For the latter, we made a uniform draw on the interval 0.0021 to 0.0028 for the base error, and calculated the total error as the square root of the square of the base error plus the square of the calibration curve uncertainty. For the simulation with 1000 samples, the first 100 samples are identical to those from the simulation with 100 samples and, similarly, for the simulation with 10,000 samples the first 100 are identical to those from the simulation with 100 samples and the first 1000 are identical to those from the the simulation with 1000 samples. For the simulation with 10,000 samples, this yielded overall measurement uncertainties ranging from 18.9 years to 26.5 years, with a mean of 22.5 years.

The simulated radiocarbon measurements were used as inputs to our algorithm and also to calculate SPDs for comparison (see Methods). We used the Hamiltonian MCMC algorithm implemented in the Stan programming language to make a total of 10,000 samples (four chains with 2000 warmup samples and 2500 non-warmup samples) from the posterior of a two-component truncated Gaussian mixture. For each such sample, the density was calculated as a function of calendar date. In turn, for a given calendar date, the quantiles were numerically determined using the 10,000 posterior samples from the HMCMC sampling. This is the basis for the posterior curves plotted in red. The 50% quantile (median) provides a single summary estimate of the density, whereas the 2.5% and 97.5% quantiles summarize the plausible range of densities for each calendar date. As *N* gets large, the SPD asymptotically approaches a stable distribution that is heavily influenced by the shape of the radiocarbon calibration curve and is not the target distribution, while our algorithm asymptotically converges to the target density.

### Identifiability of Gaussian Mixture Models

Although the simulation results in the preceding section suggest that, with sufficient observations, it is possible to precisely estimate the true parameter vector in the simulation, they are merely that: suggestive. In this section we provide evidence that this is in fact true. Specifically, we demonstrate that the two-component Gaussian mixture parameterization used in the simulations satisfies *local identifiability* and, in addition, numerically test a large number of pairs of parameterizations to demonstrate that none are observationally equivalent, which suggests that the Gaussian mixture model is *globally identifiable*.

Fundamentally, the question we address here is: can two distinct hypotheses – that is, distinct parameterizations of the Gaussian mixture model – be distinguished from each other given enough observations? This question of identifiability is a major topic in statistics, economics, and a number of other fields^17–21^. In archaeology, the related, concept equifinality has merited discussion albeit as a theoretically reverse construct from identifiability (i.e., two models are equifinal if they yield indistinguishable models)^22, 23^. More formally, an identifiability problem exists if two distinct parameterizations of a model – equivalent to two different hypotheses – yield the same probability distribution function for observed measurements. If this is the case, those distinct parameterizations are said to be observationally equivalent since there is no way to disambiguate them from data. A particular point in parameter space is identifiable if there is no other point in the parameter space allowed by the model that is observationally equivalent. If this condition is met, the model is said to be globally identifiable at that point in parameter space. A less restrictive form of identifiability – which also applies to a point in parameter space and is typically more easily demonstrated – is local identifiability. A point is locally identifiable if there is no nearby point that is observationally equivalent. Therefore, a model is globally/locally identifiable if all valid points are globally/locally identifiable^18^.

These definitions are particularly useful for the analytical demonstration of identifiability in continuous parametric models. Because the radiocarbon calibration curve is defined at discrete points (although the methodology used to determine the INTCAL, SHCAL and MARINE calibration curves in principle supports a continuous representation^24^), an analytic treatment is precluded. We can nonetheless assess identifiability numerically. To do so, we use a vector of density function values evaluated at a discrete set of points and linearize the dependence of this density vector on the underlying parameter vector to show that the null space of the matrix linking these two vectors has a dimension of zero.

Applying this approach, we show that for the time period used in our simulations, two-component Gaussian mixtures are identifiable from sets of radiocarbon dates, despite measurement uncertainty and substantial equifinality caused by the radiocarbon calibration curve. Full details are available in the supplementary information. We tested 100,000 randomly drawn parameterizations using the simulation prior probability for local identifiability (along with the actual parameterization used in Figure 1). All but one were locally identifiable, and the single sample that failed did not fail due to radiocarbon equifinality but rather because the randomly determined means and standard deviations of the two mixtures were nearly identical. We also tested 100,000 random pairs of parameter vectors to numerically determine whether any pairs were observationally equivalent. None were, and all 200,000 parameter vectors satisfied local identifiability.

## Maya Lowlands

### The MesoRAD ^14^C dataset

Aside from establishing the statistical and theoretical appeal of our end-to-end Bayesian approach, as outlined above, we also demonstrate how our approach supports sophisticated statistical inference and hypothesis testing by leveraging the Bayesian samples of the parameterized generating distribution. To do so, we utilize a large set of radiocarbon dates from the Maya lowlands and compare the results of region-wide Bayesian modeling with models that use only radiocarbon dates from the paramount Classic period polity of Tikal, located in the central Petén region of Guatemala. The dataset analyzed for this study includes radiocarbon dates from known Maya sites from across the southern and northern Maya lowlands collected from the published literature as part of the Mesoamerican Radiocarbon Database (MesoRAD^25^) (Figure 2A). In MesoRAD, associated information was recorded for each date, including the context, type of material dated, laboratory sample number, conventional ^14^C date and standard error, and the reference publication. These ^14^C dates originate from 25 environmental zones in the Maya lowlands (following^26^) that roughly correspond to the extent of shared Classic (ca. AD 250-900/1000) period ceramic, architectural, and epigraphic traditions. While MesoRAD is not a completely exhaustive dataset, it represents the largest compilation to date of published radiocarbon dates from the Maya region.

While our end-to-end approach resolves the summary problem to a large extent, it does not inherently resolve the bias problem. Therefore, we offer here an assessment of potential bias problems, which we consider relatively minor. It is useful to consider first the processes and transformations that, ultimately, link an assemblage of radiocarbon dates such as MesoRAD with living populations of the past^1, 27^. At a minimum, these include:

1. A living population generates dateable material.
2. The amount of material is proportional to population size.
3. The material gets deposited in recoverable contexts.
4. Geological and taphonomic processes filter the material that persists to the present.
5. Archaeological sites are identified via survey, construction, or other means.
6. Archaeologists choose to excavate some of those sites.
7. A subset of dateable material is in fact dated.
8. A subset of dates are published or otherwise available for meta-analysis.

**Figure 2.**
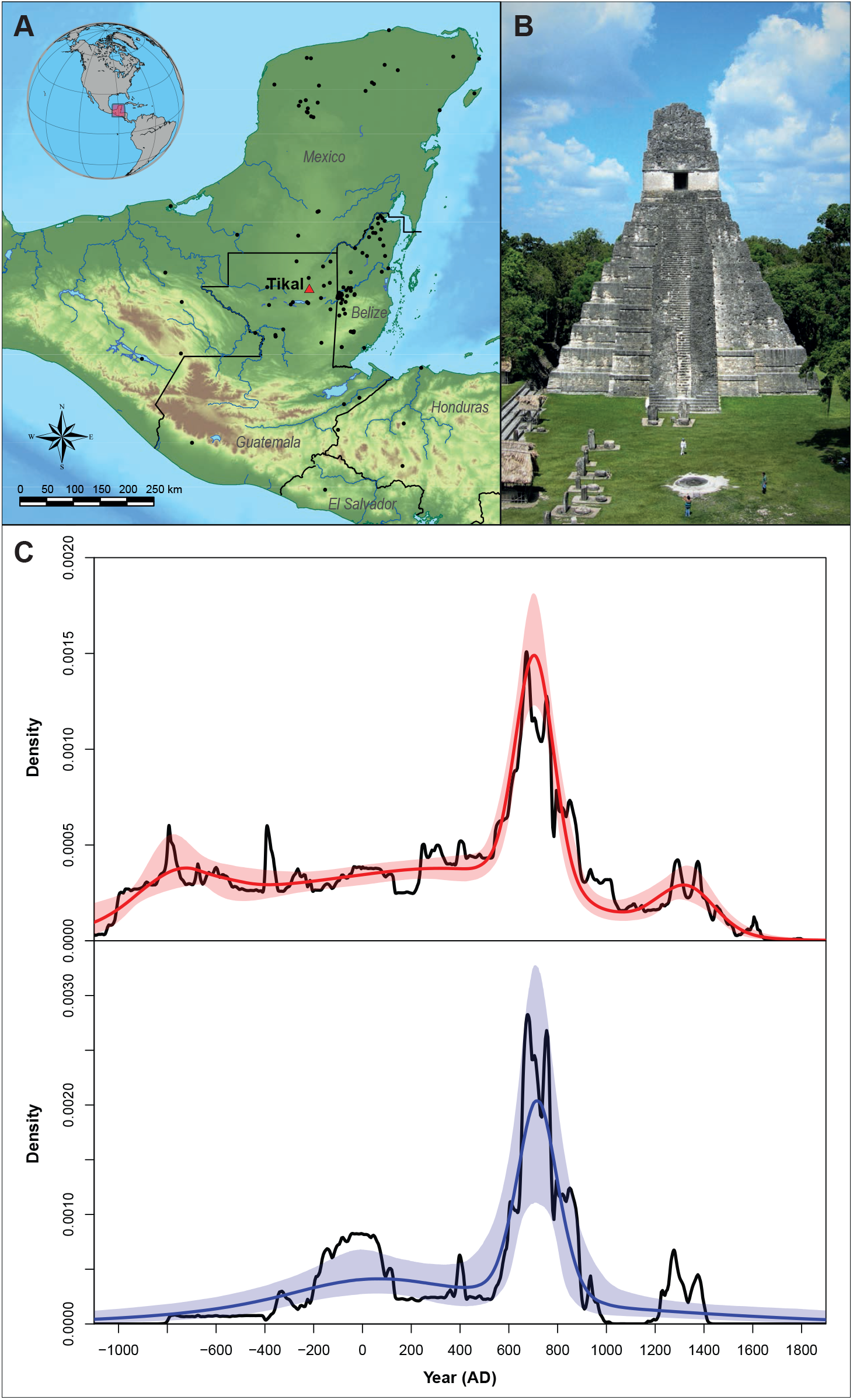
A. Map of the Maya lowlands including all sites represented in the MesoRAD dataset, B. Temple I of the Great Jaguar at Tikal, and C. Summaries from the MesoRad database including the all radiocarbon samples from the Maya lowlands (top/red) and for the site of Tikal (bottom/blue). The solid lines show the 50% quantiles (median) and the shaded areas delineate the 2.5% and 97.5% quantiles. The black curves show the respective SPDs for comparative purposes.

Following Rick^1^, we group (1-3) as Creation biases, (4) as Preservation bias, and (5-8) as Investigation biases. Approaches that have been proposed to address the bias problem include improvements in sampling strategies, explicit recommendations for cleaning datasets, implementing taphonomic corrections, and using more accurate AMS radiocarbon dates^28–30^. Regarding creation and preservation biases, cultural practices and technology are sufficiently constant in the Maya lowlands and the time periods of interest (such as the Preclassic through the Postclassic) that the total generation of dateable material is likely reasonably constant across space and time. Preservation is perhaps more variable, but also reasonably constant. Regardless, and most importantly, availability of dateable material is not a constraint to archaeological excavations in the Maya region. That is, in choosing how many dates to run, archaeologists are probably most sensitive to architecture, stratigraphy, and research goals.

Regarding investigation bias, there is a perception that in the Maya lowlands both survey and excavation have focused primarily on the Classic period, and especially the Late Classic. However, continuing research has now made it clear many parts of the Maya lowlands had large Preclassic populations. These developments are best represented at the Petén sites of El Mirador and Nakbe, where an elite ruling class mobilized local labor for massive construction projects between 600-300 BC^31–33^. A limited number of dates, and their corresponding contextual information, however, have been reported for these sites^33^, though additional radiocarbon data for early contexts is increasingly available for other regions of the lowlands.

Working with earlier contexts requires increasing reliance on radiocarbon dates to build and complement other relative dating methods. In our example, approximately 46 percent of dates from the Maya lowlands reported in this study are Preclassic (1700-2950 BP). Similarly, at Tikal, 27 percent of dates are associated with Preclassic activity. Other lines of evidence, such as ceramic analysis from excavations, record Preclassic activity in various locations across the epicentral Tikal (e.g. Mundo Perdido^34^; North Acropolis^35^), indicating a strong correspondence between the ceramic and radiocarbon records.

After chronometric hygiene and accounting for duplicates and replicates (see Supplementary Information), we analyzed a set of 1097 radiocarbon dates from 113 lowland Maya sites dating between 2850 and 200 BP^25^. We also analyzed a subset of these data from the site of Tikal, for which there were 69 observations spanning 2538 to 609 BP. Tikal represents an ideal location to understand demographic changes in the Maya lowlands given its prominence as an urban center from the Late Preclassic through Terminal Classic periods (250 BC – AD 930), as well as the extensive amount of radiocarbon dating that has taken place through archaeological investigations in the site’s epicenter^36–41^ (Figure 2B). Furthermore, comprehensive settlement-pattern, bioarchaeological, and other archaeological data from more than 70 years of systematic investigations at the site^42–48^ offer critical information for framing demographic reconstructions.

We believe the resulting dataset is representative and adequate for reconstructing human demography in the Maya lowlands and Tikal. Nevertheless, it is possible that some investigation bias remains, with a relative abundance of Classic dates compared especially to Preclassic dates. This is not only a problem with the radiocarbon assemblage, however; it is a problem with the broader archaeological and historical assemblage. Crucially, our radiocarbon-based reconstruction matches very well with primarily ceramic-based reconstructions provided by archaeologists (see Figure 3). In short, we consider the bias problem relatively minor, and any qualifications attaching to our radiocarbon-based reconstruction apply to lowland Maya archaeology as a whole.

**Figure 3.**
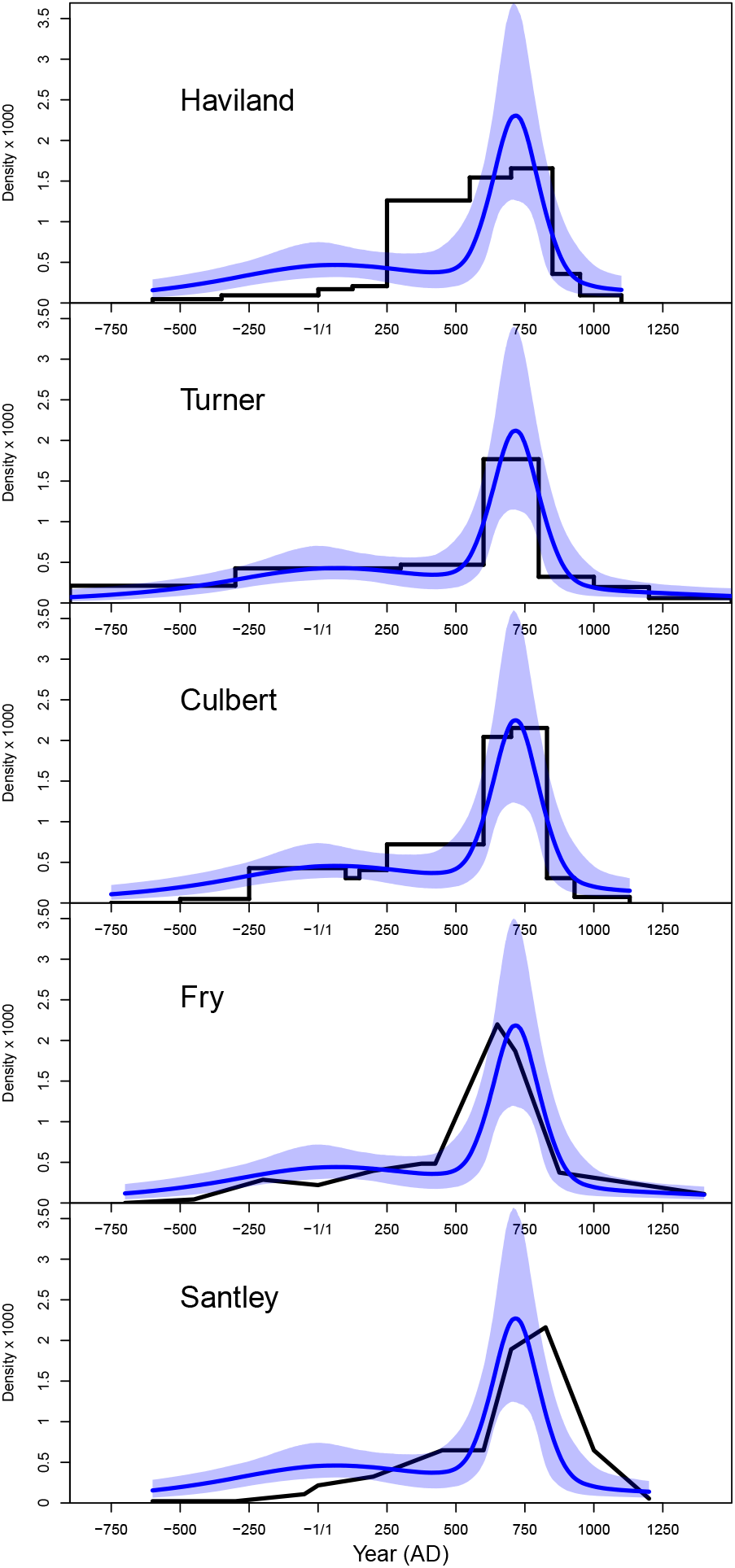
Comparison of our results as curves representing the 50% quantile (solid blue line) and 2.5% to 97.5% (shaded light blue) to five ceramic-based reconstructions of relative population change through time at Tikal (solid black lines). These estimates reflect some spatial variability, with Haviland’s (2003:Fig. 4.3)^45^ deriving from residential structures within the central 16 km^2^, Turner’s (1990:Table 15.4)^49^ from a region covering 12,600 km^2^, Culbert et al.s’ (1990:Table 5.1)^43^ estimating the relative population of small structures adjusted for 150-year average occupation within central Tikal, Fry’s (1990:Fig. 14.1)^44^ reflecting the relative population figures of central Tikal, and Santley’s (1990:Fig. 16.2)^52^ reconstructing demographic trends in central Tikal. Haviland, Culbert et al. and Turner’s estimates are over discrete time spans while Fry and Santley offer point estimates. Previous reconstructions of the the nature of population growth at Tikal suggest either protracted (Haviland) or rapid (Turner, Culbert et al., Fry, and Santley) demographic change. The analysis here points to rapid growth that initiated at the very end of the Early Classic (AD 500) followed by high population levels between the seventh and eighth centuries.

### Maya demographic trends

The resulting reconstruction of population change in the Maya lowlands is represented by the 50% quantile of the density functions resulting from the HMCMC Bayesian sampling and 2.5% to 97.5% quantile bands as with the simulation results (Figure 2C). Here and elsewhere we utilize negative AD calendar dates, which arise from calculating 1950 minus the before-present (BP) calibrated date. For both the reconstruction using all observations and that for Tikal only, we use *K* = 10 components in the finite Gaussian mixture model. This choice is validated by a detailed assessment in the Supplementary Information. Following initial growth during the Early Preclassic and Middle Preclassic (1200 – 300 BC), a strong period of rapid population growth is largely coincident with the Late Classic Period (AD 600 – 800) and declines during the Terminal Classic (AD 750 – 900*/*1000). This transient albeit tremendously important phase of growth and decline is even more clearly suggested by a growth-rate curve derived from the density function where both the periods of significant increase and decrease are highlighted (Figure 4). A secondary period of depopulation that coincides with the European conquest observed in the density curve is also present in the growth rate curve for the broader MesoRAD dataset. In Figure 2, we show not only the Bayesian reconstructions, but also the SPDs (black curves). For both cases (All data and Tikal), the SPDs often does not lie within the quantile bands of the Bayesian reconstructions.

**Figure 4.**
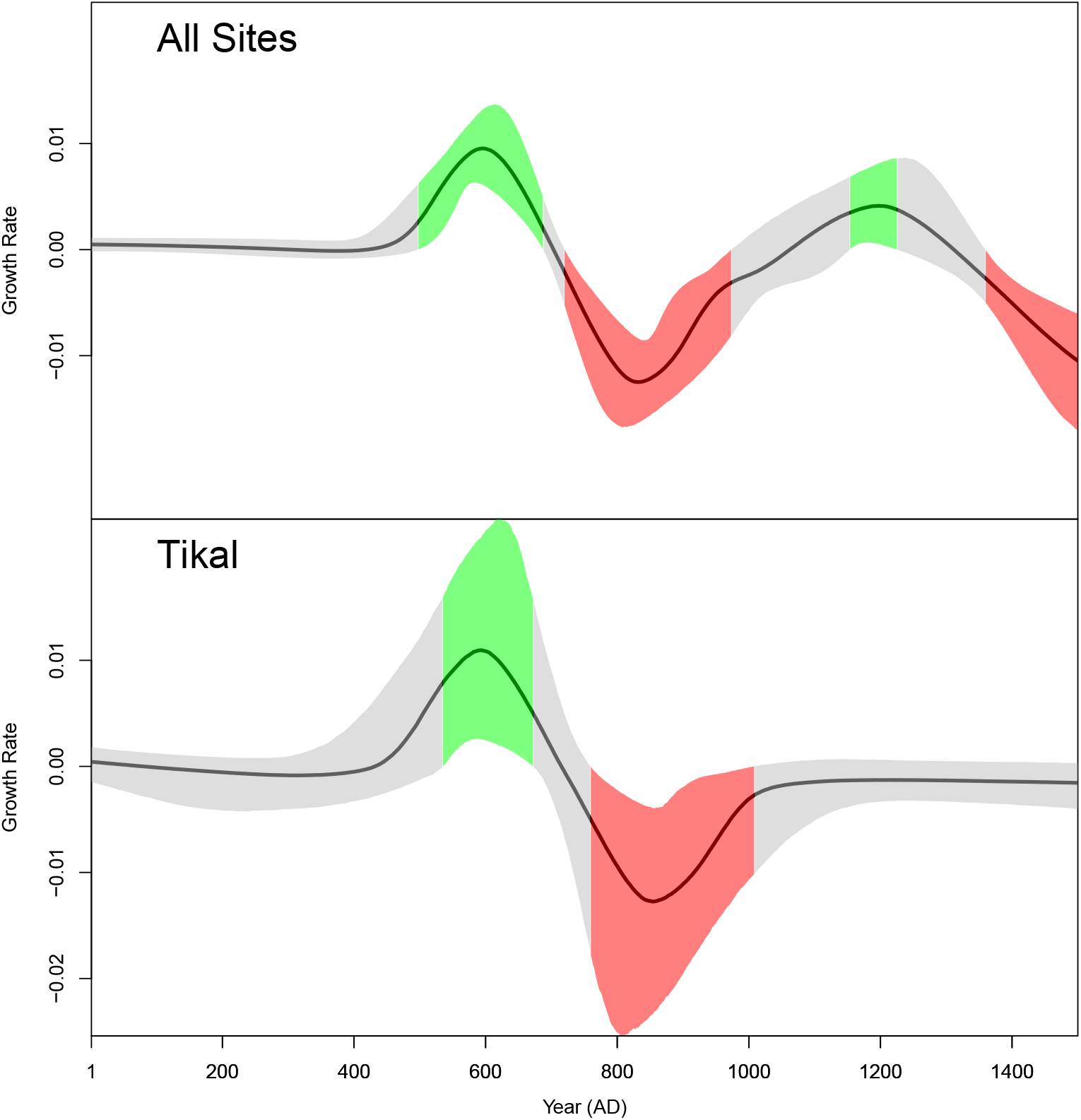
Quantile plot of per annum growth rates for all lowland Maya radiocarbon data (top) and Tikal (bottom). The solid black curves show the 50% quantiles, and the solid regions delineate the 2.5% to 97.5% bands. Green indicates regions where the 2.5% quantiles are greater than zero and red indicates regions where the 97.5% quantiles are less than zero.

The prominent center of Tikal has been the focus of extensive demographic reconstructions by archaeologists, given its importance in the Classic Maya political landscape. Previous demographic reconstructions for the site are based on ceramic chronologies from tested settlement groups^35, 43–45^. The results of the end-to-end modeling show strong parallels with the demographic reconstructions from settlement histories, which suggest low-level occupation (< 25%,^44^) in central Tikal from 750 BC to AD 515, followed by substantial population growth. Given the large temporal span of the ceramic complexes, different scholars have suggested variable timing for the initiation of substantial population growth, as well as its peak. Haviland^45^ supports an extended period of growth that began in the Early Classic period (AD 250–550), when occupation increased from around 10% to 65%, with the highest population levels (65–90%) maintained into the Late Classic (AD 550-850). Other demographic reconstructions support interpretations of rapid growth. Culbert et al.’s^43^ relative adjusted population of small structures in central Tikal shows high population growth from the Early Classic Manik complex (33.5%) to the Late Classic Ik complex (94.9%). Turner^49^ argued that Tikal’s Early Classic (AD 300–600) population was around 26.6%, increasing to 100% between AD 600–800. Fry’s reconstruction^44^ places the initiation of rapid population growth at Tikal in the later portion of the Manik phase (AD 250–600). Webster^48^ refutes interpretations of protracted growth, arguing for rapid growth that was constrained within the Late Classic Imix ceramic complex (AD 700–870). Our results support Turner’s^49^ demographic reconstruction, also closely aligning with Culbert et al^43^. It also affirms Webster’s synthetic assessment: population growth at Tikal was quite rapid and not protracted, with substantial growth initiating around AD 500 and high population levels (> 75%) between AD 660–770. In addition, our demographic reconstruction identified significant decline (> −1%, Figure 4) with populations reaching Preclassic to Early Classic levels (< 25%) by the turn of the tenth century. As identified in the settlement data, we found that the site did not experience a significant occupation during the Postclassic period as has been recorded for some areas such as in the nearby Petén Lakes^50, 51^.

### Statistical inference on Bayesian samples

Figure 5 shows histograms for three measures derived from the Bayesian samples: the year of peak population, the growth rate in AD 600, and the ratio of Late to Early Classic mean population. Each Bayesian sample represents an alternative population curve, and for each curve one can calculate the salient measure. For example, the upper plot in Figure 5 shows the distributions of the calendar date of the peak population for the two analyses (all data from the Maya lowlands, red, and Tikal only, blue). While the Tikal analysis yields a slightly later mode for the peak population distribution than for the entire dataset, there is substantial overlap in the distributions. The wider distribution for Tikal in all of the histogram plots arises from the smaller sample size for Tikal (69 versus 1097), which results in a greater range in parameter space across Bayesian samples. Archaeologists’ most recent temporal estimates for the population peak at Tikal have been broadly placed in the Ik and Imix complexes, between AD 600–830^43^, AD 600–700^44^, AD 700^45, 49^, and a bit after AD 700^48^. The end-to-end Bayesian analysis yields a peak population date at Tikal around cal AD 725, most closely in alignment with Webster’s^48^ interpretations but also within the range of many of the proposed peak dates of AD 700. Additionally, while previous demographic reconstructions have been restricted to a temporal resolution of several hundred years based on relative dating of the associated ceramic complexes, the Bayesian demographic reconstruction shows that the low-level population growth that was typical of Preclassic to Early Classic Tikal increased rapidly around AD 500 to its peak, in a span of approximately 200 years.

**Figure 5.**
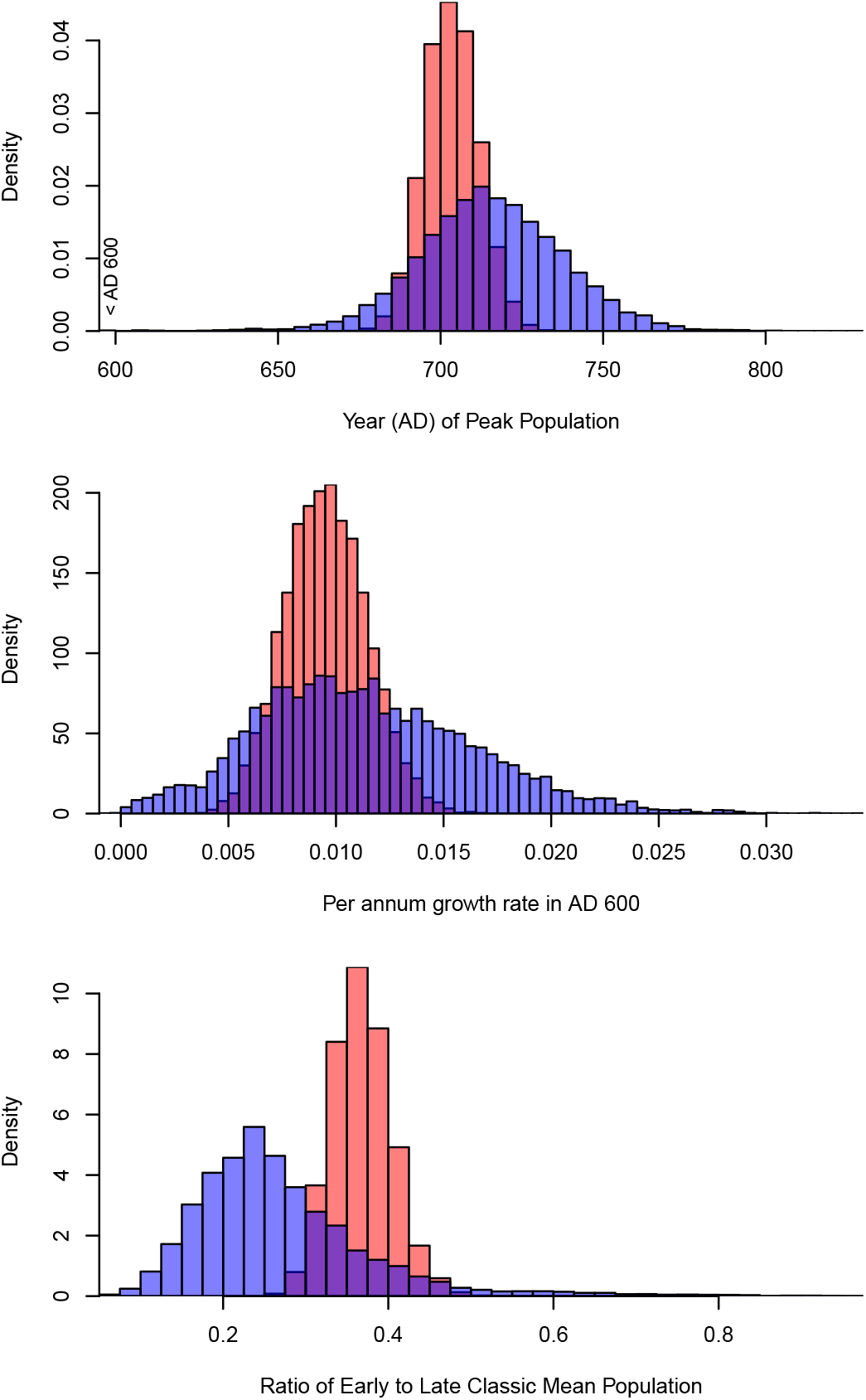
Distributions of a particular variable derived from the Bayesian samples, where each Bayesian sample parameterizes a separate population curve. The upper plot shows the calendar date of the peak population. The middle plot shows the per annum growth rate in AD 600. The bottom plot shows the ratio of the mean population in the Late Classic (AD 550–830) to that in the Early Classic (AD 250–550). As in Figure 2, red is for all radiocarbon samples and blue for Tikal. For Tikal in the peak population histogram, 4 samples (out of 10,000) with values less than AD 600 have been placed in the bin AD 595–600 to improve interpretability.

The middle plot in Figure 5 shows the per annum growth rate in cal AD 600.^1^ Substantive efforts have been made to frame demographic growth rates throughout prehistory^53^, with some scholars suggesting long-term population growth rates averaging between 0.1 and 0.2% for many areas of the world^54, 55^. In Mesoamerica, periods of growth were characterized between 0.39% and 0.59% at Ixtapalapa, Mexico, whereas the annual growth rates from the highland site of Kaminaljuyu, Guatemala, were around 0.1%^56^. Turner^49^ describes the “population explosion” between AD 600–800 across the Maya Lowlands as being characterized between by 0.45-0.58% growth per annum. Santley^52^ suggests a crude growth rate for the entire central and southern Maya lowlands in AD 600 around 0.1%, increasing to around 1.4% in AD 700. This maximum estimate is close to the growth model proposed for the lowland Maya site of Copan^57^ between AD 749–799. Overall, the annual growth rate identified from radiocarbon data at Tikal in cal AD 600 falls within typical to high levels identified in cases of rapid population growth both in Mesoamerica, as well as for other cases of rapid growth around the world. It remains an open question whether this population increase resulted from natural population increase, migration, or some combination of the two.

Finally, the lower plot of Figure 5 shows the ratio of Early to Late Classic mean population at Tikal. These values suggest that the Late Classic population may have been between two to four times the size of the Early Classic population. Webster’s compilation^48^ of settlement-based population estimates for Tikal fall between 20,000 to 28,000 during the Early Classic Manik ceramic complex (AD 250–600). These population estimates increase to between 42, 000 and 58, 000 during the Late Classic Imix ceramic complex (AD 700–830). With these settlement-based estimates (with ratios between 2 to 3) in mind, the end-to-end Bayesian modeling estimates for Late Classic to Early Classic population align well with the archaeological evidence from Tikal. Although total population estimates for Tikal are debated, in part because of the recent acquisition of full-coverage lidar (light detection and ranging) remote sensing surveys^58^, the Bayesian modeling offers a generalized method of examining overall demographic changes through time. Furthermore, the end-to-end approach provides a set of population reconstructions consistent with the input set of radiocarbon calibration dates than can be used to actively characterize the uncertainty of any measure of interest, such as the population growth rate in a particular year or the ratio of mean populations across time periods.

Results from the analysis of Tikal dates demonstrate the utility of the end-to-end approach we propose by showing strong alignment with settlement-based demographic reconstructions that have suggested rapid population growth in the Late Classic period (AD 550–830). Although the nature of demographic growth, as well as the timing of the peak in population, have been highly debated, the Bayesian demographic reconstruction here points to a period of rapid population growth that peaked at the beginning of the eighth century AD. Such rapid changes in growth suggest improved infant/child survival or recruitment^59, 60^ since these are the demographic rates to which the rate of intrinsic increase are most sensitive. Infant mortality data are not available for Tikal, although Wright and White’s^61^ synthesis of osteological studies for other lowland Maya sites suggests that there were no statistically significant chronological differences in child health in the context of Late Classic population increase. Current studies, however, are limited by small bioarchaeological sample sizes.

### Limitations and future work

Migration into the urban center may also account for the rapid population growth at the site and the florescence of the city may have provided recruitment opportunities previously unavailable to commoners. Wright’s^62^ analysis of strontium and oxygen isotopes at Tikal showed the highest non-local values (27.5%) during the Early Classic period, decreasing (10.2%) during Late Classic times. Future studies may help to illuminate these vital questions. Overall, when compared to the archaeological data from more than 70 years of investigations at Tikal, the results highlight the ability of the end-to-end Bayesian approach to resolve questions that have persisted for decades.

Indeed, it would be straightforward to extend the end-to-end approach used in this article to incorporate both age-structure and population-structure into the demographic model, and thus use additional types of information such as skeletal age-at-death data and isotopic data informative of migration to further refine the demographic reconstruction. In fact, this would not only help to address the summary problem that is the focus of this article, but also to identify the extent of bias and help to alleviate this. The reason is that different types of data are subject to different biases. For example, all else being equal, in a growing population, the age-at-death distribution contains more young people, whereas in a shrinking population it contains more old people. If the pattern observed in skeletal data is inconsistent with the growth rates reconstructed from radiocarbon data, there may be a bias in one or both types of data, and fusing the data using end-to-end inference will yield a better population reconstruction.

In this article, we parameterized the generating distribution using a finite Gaussian mixture model with *K* = 10 mixtures for the analysis of Maya dates. Both the choice of a Gaussian mixture model and number of mixture components could be the subject of future work. For example, while in the Supplementary Information we validate the choice of *K* = 10 for the number of components, future analyses could instead use Bayesian model selection to the chose the number of mixtures^63^ or specify a prior over the number of mixture components.

## Conclusions

Compilations of radiocarbon dates present an important scientific source for scholars interested in reconstructing demographic change over time at multiple spatial scales. The methods used to reconstruct population histories from radiocarbon dates have known limitations. In particular, these methods suffer from both bias and summary problems. Conceptual and methodological improvements, and increased sample coverage, have increasingly helped to resolve the bias problem but the summary problem has been largely overlooked. In this paper, we presented a new approach that greatly increases the utility of inference from compilations of radiocarbon dates, greatly improving their ability to resolve population histories and thereby marking a notable advance in solving the summary problem. Compared with the current tools of choice, SPDs and KDEs, the end-to-end Bayesian approach can more effectively estimate the original distribution from which a sample is drawn by accommodating ambiguity at a level higher than the individual sample, and furthermore allows direct inference over demographic parameters such as the growth rate. The approach combines finite-mixture models and Bayesian inference. We show both numerically and analytically that the models are identifiable and ultimately converge to the generating distribution with simulated data sets. Applying the model to a database of radiocarbon samples from the Maya Lowlands, we are able to provide some critical tests of existing hypotheses about the population history of the region, and for the site of Tikal in particular. The nature of Tikal’s population growth has been extensively debated by archaeologists, and our demographic reconstruction supports an initial phase of slow population increase in the Preclassic and Early Classic, followed by a period of dramatic population increase for the century spanning cal AD 550-650 with values at the peak possibly in excess of 1% annually. We estimate the calendar data of the peak population size at Tikal at circa AD 725 and decline by AD 770. Our analysis supports the demographic reconstructions presented by Turner^49^, and verifies Webster’s^48^ assertion that Tikal’s trajectory of growth was episodic and not protracted. For the Maya lowlands as a whole, we estimate very robust growth, in excess of 1% annually, for approximately the same interval over which Tikal grew rapidly. These results are significant because they suggest that the population boom was likely linked to increased childhood health or recruitment opportunities (and perhaps also to migration). Overall, we see many opportunities to apply the tools described in this paper to the rapidly-expanding corpus of radiocarbon records and to ask increasingly sophisticated demographic questions from these data.

## Data Availability

The MesoRAD database is archived with tDAR (The Digital Archaeological Record) with the doi 10.6067/XCV8455306. The data are also available in the file MesoRAD-v.1.1_FINAL_no_locations.xlsx in the github repository eehh-stanford/price2020. Latitude/longitude locations have been removed from the publicly available data.

## Acknowledgements

We thank Darcy Bird, Brian Codding, Dan Contreras, and Chris Jazwa for helpful comments.

## Author Contributions

All authors wrote the manuscript. MHP designed and implemented the end-to-end Bayesian algorithm and identifiability analysis. KB contributed to the software packages. JAH and CEE collated the radiocarbon data (MesoRAD).

## Footnotes

1 For both Figure 4 and Figure 5, the per annum growth rate is calculated by taking the logarithm of the instantaneous growth rate (see Methods) measured in units of inverse years.

