## Supplementary Information for "End-to-end Bayesian analysis of ^14^C dates reveals new insights into lowland Maya demography"

### Supplementary information for “End-to-end Bayesian analysis of radiocarbon $^{14}C$ dates reveals new insights into lowland Maya demography” by Michael Price et al.

**3 Supplementary Note 3: Choice of number of mixtures for Maya analysis** **9**
**Supplementary References** **12**

#### 1 Supplementary Note 1: Identifiability of Radiocarbon Measurements

Under what conditions can two distinct hypotheses be distinguished from each other? This question of identifiability is a major topic in statistics, economics, and other fields. Conceptually, an identifiability problem exists if two distinct parameterizations of a model yield the same distribution function for observed measurements. If this is the case, those distinct parameterizations are said to be observationally equivalent. A particular point in parameter space is identifiable if there is no other point in parameter space allowed by the model that is observationally equivalent. This is global identifiability. A less restrictive form of identifiability – which is also typically more easily demonstrated – is local identifiability. A point is locally identifiable if there is no nearby point that is observationally equivalent. For a more formal and extended treatment of these concepts and terms, see[1].

In this section, we adopt the conceptual framework described in[1] and in the citations in the main text, but utilize methods from linear algebra that are appropriate given the discrete nature of the calibration curve and the discrete Riemman approximation we utilize to calculate the likelihood function for a set of radiocarbon observations. We adopt the variable definitions and notation in the Materials and Methods of the main text: $\theta$ parameterizes the density $(p(t|\theta))$, $\tau$ (indexed by $g$) is the set of grid points for Riemann integration, $\mathbf{M}$ is the measurement matrix with elements $M_{ig}$ ($i$ indexes locations where the fraction modern, $\phi$, is evaluated), $v_g(\theta) = p(\tau_g|\theta)$ defines the sampling vector $\mathbf{v}$, and $\mathbf{h} = \mathbf{M}\mathbf{v}$ is the vector of likelihoods (i.e., the probability density of the observed fraction modern). Since the locations at which $\phi$ is evaluated in this supplement no longer necessarily correpsond to measurements, we drop the $m$ subscript, $\phi_{m,i} \rightarrow \phi_i$. We offer three definitions:

**Definition 1** *Two parameterizations $\theta_1$ and $\theta_2$ are observationally equivalent if $\mathbf{h}(\theta_1) = \mathbf{h}(\theta_2)$.*

**Definition 2** *A parameterization $\theta_0$ is identifiable if there is no other $\theta$ that is observationally equivalent.*

**Definition 3** *A parameterization $\theta_0$ is locally identifiable if there exists an open neighborhood around $\theta_0$ containing no other $\theta$ that is observationally equivalent.*

Let $\mathbf{h}_1 = \mathbf{h}(\theta_1)$ and $\mathbf{h}_2 = \mathbf{h}(\theta_2)$. Setting these vectors equal to each other, $\mathbf{h}_1 = \mathbf{h}_2$, yields

$$\mathbf{M}(\mathbf{v}_2 - \mathbf{v}_1) = \mathbf{0}, \tag{1}$$

where $\mathbf{v}_1 = \mathbf{v}(\theta_1)$ and $\mathbf{v}_2 = \mathbf{v}(\theta_2)$. A vector $\mathbf{x}$ in the (right) null space of $\mathbf{M}$ satisfies

$$\mathbf{M}\mathbf{x} = \mathbf{0}. \tag{2}$$

If the null space is empty (i.e., its dimension is zero and excluding the trivial solution $\mathbf{x} = \mathbf{0}$[1]) there is no identifiability problem since $\mathbf{v}_1 \neq \mathbf{v}_2$ necessarily implies $\mathbf{h}_1 \neq \mathbf{h}_2$. Although it is not typically true that the null space of $\mathbf{M}$ is empty, it is possible that a set of dates is known to come from a portion of the radiocarbon calibration curve where the fraction modern uniquely determines the calendar date (see below). Even if the null space of $\mathbf{M}$ is not empty, there may be no identifiability problem. In particular, an identifiability problem (global or local) exists only if the difference $\mathbf{v}_2 - \mathbf{v}_2$ lies in the null space of $\mathbf{M}$.

Local identifiability can be assessed by considering a Taylor approximation to an element of $\mathbf{v}$ around the point $\theta_0$,

$$v_g \approx p(y_g|\theta_0) + \sum_j^J \left.\frac{\partial p(y|\theta)}{\partial \theta^{(j)}}\right|_{y_g,\theta_0} \Delta\theta^{(j)}, \tag{3}$$

where $\theta^{(j)}$ is the $j$-th element of the parameterization and $J$ is the total number of parameters. Utilizing this Taylor expansion yields

$$\Delta\mathbf{h} = \mathbf{M}\mathbf{P}\Delta\theta, \tag{4}$$

where the elements of $\mathbf{P}$ are

$$P_{gj} = \left.\frac{\partial p(t|\theta)}{\partial \theta^{(j)}}\right|_{t_g,\theta_0}. \tag{5}$$

The model is locally identifiable at $\theta_0$ if the null space of $\mathbf{M}\mathbf{P}$ is empty. We shall call $\mathbf{P}$ the perturbation matrix. For the likelihood calculation described in the Materials and Methods of the main text, the measurement matrix $\mathbf{M}$ is evaluated at the known fraction modern values of the samples (and utilizing the known uncertainties) and at a grid of calendar dates $\tau$. For the identifiability analysis, it is evaluated at a grid of fraction modern values and at a grid of calendar dates. We use a fixed value for the measurement uncertainty (this is varied in the exponential example, and set to 0.001 in the Gaussian mixture example). For the grid of fraction modern values, we first determine the range of fraction modern values of the calibration curve for the grid of calendar dates, then extend this range by four times the measurement uncertainty on both the low and high end. This establishes the range. We sample this range evenly using four times the length of (i.e., number of elements in) the calendar date grid.

[1] To ensure that the trivial solution is not relevant, we assume that $p(t|\theta)$ is parameterized in such a way that $\theta$ uniquely specifies the probability density function and vice versa. That is, no identifiability problem is "built in" by the way the probability density function is defined. This is necessary since certain common parameterizations do have an identifiability problem (for example, mixtures of Gaussians must be ordered by their means or otherwise modified to remove an identifiability problem that arises from swapping all the parameters for two different components of the mixture).

#### 1.1 Identifiability of exponentials

It is useful, before considering Gaussian mixtures, to consider first the identifiability of exponential growth or decay. This is a good starting point because exponentials are defined by one parameter, the growth rate $r$, and it is thus possible to build some intuition prior to considering Gaussian mixtures. Aside from this, exponentials are a commonly adopted parameterization in the dates-as-data literature[2,3], and will be of inherent interest to some readers. We assume that observations are restricted to the interval $\tau_{min}$ to $\tau_{max}$. Since the density function must integrate to 1 on this interval, the full parameterization is

$$p(t|r) = \frac{re^{rt}}{e^{r\tau_{\max}} - e^{r\tau_{\min}}}. \tag{6}$$

This definition works for both positive ($r > 0$) and negative ($r < 0$) growth. Calculation of the perturbation matrix requires taking only the partial derivative with respect to $r$,

$$\frac{\partial p(t|r)}{\partial r} = p(t|r)[\frac{1}{r} + t - \tau_{\max} - \frac{(\tau_{\max} - \tau_{\min})\exp(-r(\tau_{\max} - \tau_{\min}))}{1 - \exp(-r(\tau_{\max} - \tau_{\min}))}], \tag{7}$$

where we have subsumed the dependence of $p(t|r)$ on $\tau_{\min}$ and $\tau_{\max}$ since they are fixed (this assumption could be relaxed). For the special case of $r = 0$ one can apply L'Hôpital's rule to show that Equation 7 becomes $0.5 + (t - \tau_{\max})/(\tau_{\max} - \tau_{\min})$. Figure 6 plots the radiocarbon calibration curve on the top row and the probability density of the fraction modern, varying three things: (a) the time span, (b) the measurement uncertainty, and (c) the growth rate. The three time spans are AD 700-950 (Span 1), AD 800-850 (Span 2), and AD 900-950 (Span 3); these are delineated by red bars on the top row. The two settings for the measurement uncertainty (fraction modern) are 0.01, used for the middle set of plots, and 0.001, used for the bottom set of plots. These correspond to a radiocarbon year uncertainty (the most common way of reporting uncertainties) of 9.4 / 93.9 years in AD 700 and 9.1 / 91.0 years in AD 950. The measurement uncertainty and calibration curve uncertainty combine to yield the total uncertainty (see main text Methods and Materials). The yearly growth factors used are -4% to +4%, with a spacing of 1%; to be precise, for a yearly growth factor of 3%, $\delta = 0.03$, the growth rate is $r = \log(1+\delta) = 0.0296$, where $r$ has units of inverse years. Each of the nine growth factors yields a distinct density function for the generating distribution, $p(t|r)$, and, in turn, a distinct density function for the fraction modern, $h(\phi|r)$. The black curves in the middle and bottom row of plots are the density functions for the fraction modern, $h(\phi|r)$.

For each of the six sub-plots on the middle and bottom rows of Figure 1, an identifiability problem would exist if any two values of $r$ yielded the same density function for the fraction modern (black curves). For each of the examples (the six plots in the bottom two rows of Figure 1), we have numerically assessed the local identifiability of all 9 settings of $r$. In all cases, we confirmed local identifiability. Furthermore, it appears that for all the examples the density functions ,$h(\phi|r)$, vary smoothly with $r$, implying global identifiability, though we realize this constitutes neither a quantitative numerical check nor a proof. Perhaps more importantly, however, for some of the examples – most notably the very middle plot – it would take a very large number of observations to distinguish two different parameterizations. That being said, we deliberately chose the two smaller time spans to be vexing: both Span 2 and Span 3 lie entirely in non-ivertible regions of the calibration curve, and the calibration curve is close to flat within Span 2. Even so, the exponential parameterization is locally identifiable.

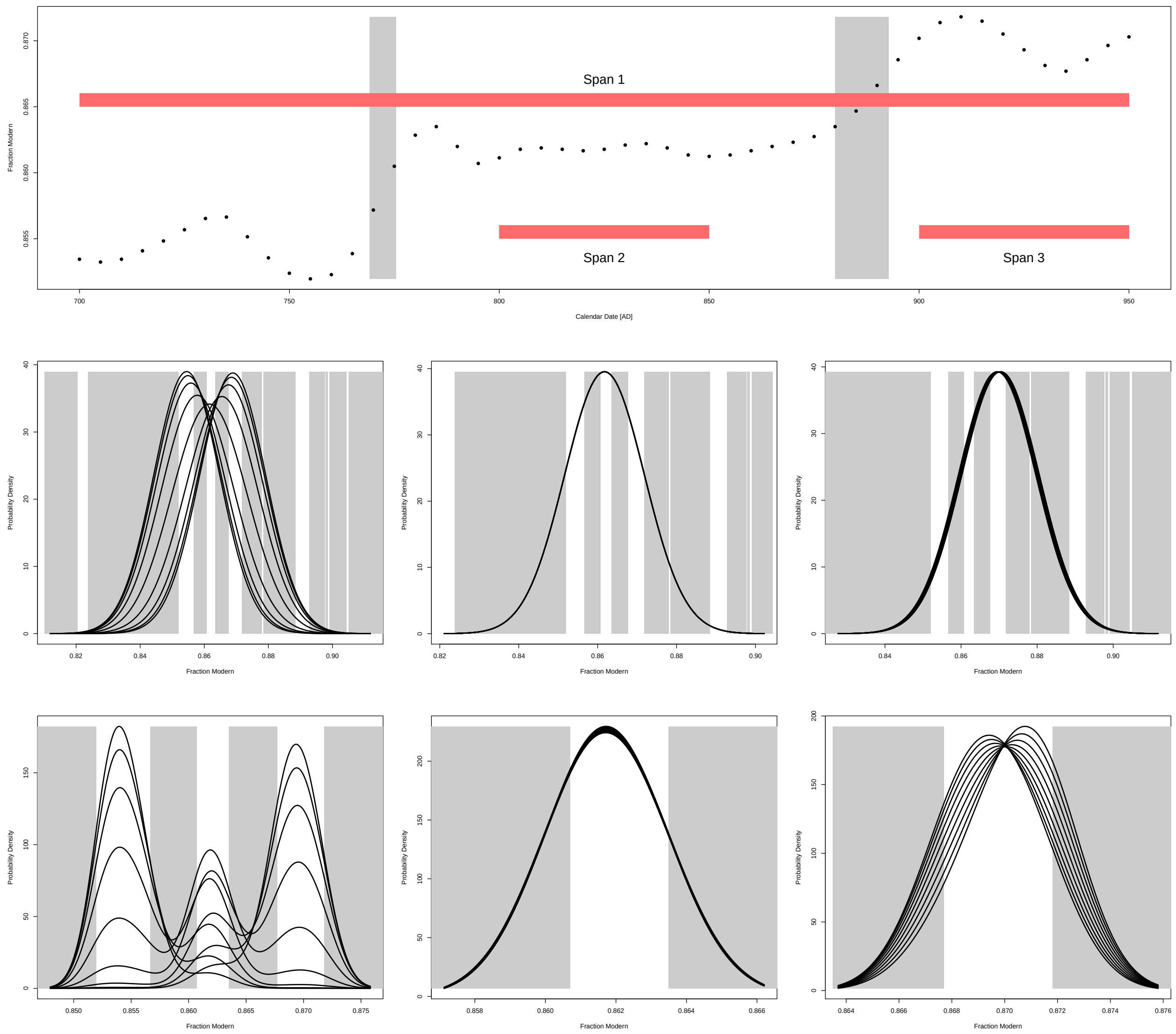

**Supplementary Figure 1.** Calibration curve (top row) and probability density functions for the fraction modern (middle and bottom rows) for a range of examples of exponential growth/decay. In the middle and bottom rows, the first column corresponds to Span 1 in the top row, the middle column to Span 2, and the right column to Span 3. The middle row utilizes a measurement uncertainty for the fraction modern of 0.01 and the bottom row an uncertainty of 0.001. The gray bands mark regions where the calibration curve is invertible (that is, there is a one-to-one mapping between calendar year and the fraction modern). Each plot in the bottom two rows has nine black curves, and each black curve in a plot corresponds to a distinct parameterization of the exponential; in turn, each parameterization of the exponential yields a distinct probability density function for the fraction modern measurements. The nine growth factors range from -4% to +4%, yearly, with a spacing of 1%. Both Span 2 and Span 3 lie completely in non-invertible regions of the calibration curve, but differ in that Span 2 is nearly flat. In all cases, there is no problem with identifiability, though in some cases the curves are close to identical, which implies that a large number of samples would be needed to distinguish alternative growth rates if, say, only data on the interval 800 to 850 were available.

#### 1.2 Identifiability of Gaussian mixtures

Gaussian mixtures are a common and flexible parameterization for probability density functions. The probability density function for a Gaussian mixture with $K$ components is

$$p(t|\theta) = \frac{1}{\eta} \sum_{k=1}^{K} \pi_k f_{\mathcal{N}}(t;\mu_k,s_k) = \frac{z}{\eta}, \tag{8}$$

where $f_{\mathcal{N}}$ is the Gaussian probability density function, we define $z$ as the unnormalized density for convenience, and $\pi_k$, $\mu_k$, and $s_k$ are respectively the mixture proportion, mean, and standard deviation for component $k$. $\eta$ is a normalization term to account for the possible truncation of each mixture at a lower and upper calendar date,

$$\eta = \sum_{k=1}^{K} \pi_k \left[F_{\mathcal{N}}(\tau_{\max};\mu_k,s_k) - F_{\mathcal{N}}(\tau_{\min};\mu_k,s_k)\right] = \sum_{k=1}^{K} \pi_k \left[F_{\mathcal{N}}(\frac{\tau_{\max}-\mu_k}{s_k}) - F_{\mathcal{N}}(\frac{\tau_{\min}-\mu_k}{s_k})\right], \tag{9}$$

where $F_{\mathcal{N}}$ is the cumulative Gaussian distribution function. We adopt the convention that if $\mu_k$ and $s_k$ are omitted from either $f_{\mathcal{N}}$ or $F_{\mathcal{N}}$ they are presumed to equal 0 and 1 respectively – e.g., $F_{\mathcal{N}}(t) = F_{\mathcal{N}}(t;0,1)$, the cumulative distribution function of the standard normal. Since the mixture components must sum to 1, we omit $\pi_1$ from the perturbation matrix and must account for the constraint $\pi_1 = 1 - \sum_{k=2}^{K} \pi_k$ in calculating the elements of the perturbation matrix. The partial derivatives satisfy

$$\frac{\partial p(t|\theta)}{\partial \pi_k} = \frac{1}{\eta}\frac{\partial z}{\partial \pi_k} - \frac{1}{\eta^2}\frac{\partial \eta}{\partial \pi_k}, \tag{10}$$

$$\frac{\partial p(t|\theta)}{\partial \mu_k} = \frac{1}{\eta}\frac{\partial z}{\partial \mu_k} - \frac{1}{\eta^2}\frac{\partial \eta}{\partial \mu_k}, \tag{11}$$

and

$$\frac{\partial p(t|\theta)}{\partial s_k} = \frac{1}{\eta}\frac{\partial z}{\partial s_k} - \frac{1}{\eta^2}\frac{\partial \eta}{\partial s_k}. \tag{12}$$

The partial derivatives of $z$ are

$$\frac{\partial z}{\partial \pi_k} = f_{\mathcal{N}}(t|\mu_k,s_k) - f_{\mathcal{N}}(t|\mu_1,s_1), \tag{13}$$

$$\frac{\partial z}{\partial \mu_k} = \pi_k \frac{t-\mu_k}{s_k^2} f_{\mathcal{N}}(t|\mu_k,s_k), \tag{14}$$

and

$$\frac{\partial z}{\partial s_k} = \pi_k \left[-\frac{1}{s_k} + \frac{(t-\mu_k)^2}{s_k^3}\right] f_{\mathcal{N}}(t|\mu_k,s_k). \tag{15}$$

The partial derivatives of $\eta$ are

$$\frac{\partial \eta}{\partial \pi_k} = F_{\mathcal{N}}(\frac{\tau_{\max} - \mu_k}{s_k}) - F_{\mathcal{N}}(\frac{\tau_{\min} - \mu_k}{s_k}) - F_{\mathcal{N}}(\frac{\tau_{\max} - \mu_1}{s_1}) + F_{\mathcal{N}}(\frac{\tau_{\min} - \mu_1}{s_1}), \tag{16}$$

$$\frac{\partial \eta}{\partial \mu_k} = -\frac{\pi_k}{s_k}\left[f_{\mathcal{N}}(\frac{\tau_{\max} - \mu_k}{s_k}) - f_{\mathcal{N}}(\frac{\tau_{\min} - \mu_k}{s_k})\right], \tag{17}$$

and

$$\frac{\partial \eta}{\partial s_k} = -\frac{\pi_k}{s_k^2}\left[f_{\mathcal{N}}(\frac{\tau_{\max} - \mu_k}{s_k})(\tau_{\max} - \mu_k) - f_{\mathcal{N}}(\frac{\tau_{\min} - \mu_k}{s_k})(\tau_{\min} - \mu_k)\right]. \tag{18}$$

The preceding equations allow calculation of the elements of the perturbation matrix. We drew 100,000 random samples of the parameter vector using $\alpha_d = 1$, $\alpha_s = 10$, and $\alpha_r = (10-1)/50$ (see main text Material and Methods for definitions) then checked for local identifiability of each by ensuring that the null space of **MP** is empty. Exactly one random draw failed this test with a null space dimension of 1. However, this failure was a result of the parameterization itself. The parameter vector components were $\pi_1 = 0.3694483$, $\pi_2 = 0.6305517$, $\mu_1 = 750.3344293$, $\mu_2 = 747.7518300$, $s_1 = 72.5960876$, and $s_2 = 72.6153552$. These have nearly identical means and standard deviations. For this case, the matrix **P** also has a null space dimension of 1 with the grid spacing used for the calculation, indicating that given the fidelity of the overall test it is the means and standard deviations being equal that caused the lack of local identifiability.

We also randomly sampled 100,000 pairs of parameter vectors to assess whether they yielded identical probability density functions for the fraction modern. We used the same measurement matrix and sampling parameters as for the preceding check of local identifiability. We considered a pair to have the same density if the difference in densities never exceeded $1e-6$ times the mean value of the two density functions. No pair failed this test. We further checked each draw from the pair for local identifiability, and none failed local identifiability. Overall, therefore, we checked 300,000 parameterizations for local identifiability, with none failing due to equifinality of the radiocarbon calibration curve.

To provide some intuition for these checks, Figure 2 (middle plot) shows 8 two-component Gaussian mixture samples defined on the interval AD 600 to 1300. All 8 curves are locally identifiable. The top plot in Figure 2 show the calibration curve. As in other figures, the gray bands delineate invertible regions of the calibration curve. The bottom plot shows the probability density function over radiocarbon measurements (fraction modern) for each parameterization assuming a measurement uncertainty of the radiocarbon determinations of 0.001, which corresponds to an uncertainty of 9.2 uncalibrated years at the central date of 950. The gray bands in the bottom plot delineate invertible regions of the calibration curve defined over the fraction modern (as opposed to over calendar dates as in the other two plots in Figure 2).

Radiocarbon measurements tend to cluster in the non-invertible regions. For example, consider the time span AD 687.9 to 880, which fully contains two non-invertible regions and is highlighted by a horizontal red bar in the middle plot of Figure 2. A horizontal red bar also highlights the corresponding span of fraction modern values in the bottom plot of Figure 2. The first non-invertible region spans AD 687.9 to 769.2 for calendar dates and 0.852 to 0.857 for fraction modern, whereas the second non-invertible region spans AD 775.4 to 880 for calendar date and 0.861 t0 0.863 for fraction modern. Since the calibration curve is fairly flat in these regions, the radiocarbon measurements from samples originating at the calendar dates in this time span are close to each other, which leads to the clustering of observations in the bottom plot of Figure 2. Nevertheless, all the curves are distinct, so there is no problem of identifiability.

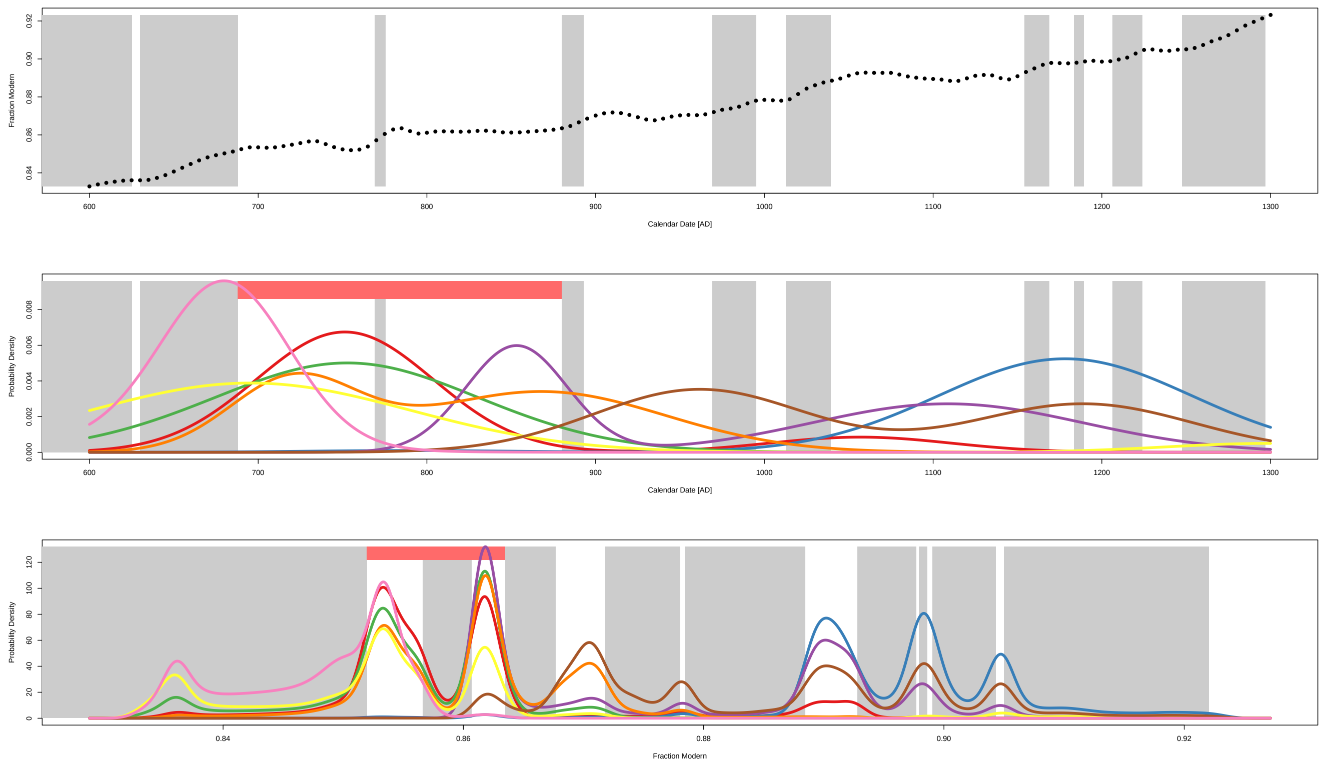

**Supplementary Figure 2.** Gaussian mixture example

#### 2 Supplementary Note 2: Lowland Maya 14C Dataset Details

A total of 1529 radiocarbon dates from the Maya lowlands were collected from the published literature as part of the MesoRAD database. The database includes radiocarbon dates and isotopic data from archaeological sites in across Mesoamerica. The database is currently archived with tDAR (The Digital Archaeological Record; doi:10.6067/XCV8455306).

In addition to being available on tDAR, the file MesoRAD-v.1.1_FINAL_no_locations.xlsx in the github repository eehh-stanford/price2020 shows radiocarbon dates organized by uncalibrated calendar date, along with region and site. Associated information was recorded for each date, including contextual information (specific stratigraphic and spatial relationships reported), type of material dated (e.g., charcoal, human remains, faunal remains), lab number, conventional 14C date and error ranges, 2-$\sigma$ calibrated distributions, whether the sample was dated via Accelerated Mass Spectrometer (AMS) or conventional 14C dating, and reference publications.

Dates in from the Mesoamerican Radiocarbon (MesoRAD) database were subjected to chronometric hygiene criteria established by Hoggarth and colleagues[4]), page 31, to eliminate questionable dates from the sample prior to modeling. In[4], dates were categorized according to the following criteria:

1. The association with cultural remains cannot be ambiguous and the date should be supported by additional archaeological evidence. A lack of contextual information or lack of reporting for uncalibrated conventional dates yields ambiguous associations.

2. Dates from bone (human or faunal) and/or shell require additional pretreatment, purification, and assessment of reservoir effects or other environmental corrections. When not corrected for diagenesis or reservoir effects, calibrated date ranges can potentially be erroneous.

3. Only radiocarbon dates with measurement precisions below ±100 14C years should be considered for modeling, unless they can be constrained through stratigraphic modeling, since larger errors contribute to "blurred probability distributions" and impede clear chronological distinctions (see also[5]).

4. Dates derived from experimental techniques that have yielded questionable results in the region should be rejected until proven more reliable.

MesoRAD-v.1.1_FINAL_no_locations.xls includes a description of reasons of chronometric hygiene where applicable. Not all notes in the column constitute an absolute reason for rejection for "dates as data approaches", though some issues associated with dates not in alignment with cultural remains (chronometric hygiene criteria 1) or outliers in sequences would cause enough concern for rejection. Researchers using these data for future studies should use their own judgement, based on the methods that are used. Though we have previously rejected dates with measurement precisions above ±100 14C years, the End-to-End Bayesian modeling can deal with these large distributions and therefore we have retained those for this analysis. For the purposes of this study, dates were rejected when no conventional 14C year was reported or when it was too early or too late for the context as reported by the original investigators.

In addition to the information provided above, the MesoRAD database provides information on which radiocarbon determinations are duplicates or replicates. Duplicates are dates recorded from the same object (e.g., burial, wooden lintel, piece of charcoal) and are used to provide information on the error statistics of the overall radiocarbon measurement process. Replicate dates, on the other hand, are those that derive from the same context, but are not necessarily from the same material or preparation chain. Replicate samples are used, for example, to assess the possibility of erroneous dates from old wood[6].

We chose to treat both duplicates and replicates as representing only a single event contributing to the overall generating probability distribution. Therefore, for each set of duplicates/replicates we used the combination_Gauss function in the ArchaeoChron R package[7] to create a pooled estimate from each set of duplicates/replicates with the pooled mean and error in uncalibrated radiocarbon years before present.

#### 2.1 Tikal Radiocarbon Data

Over 117 radiocarbon dates from the large polity of Tikal have been published, all of which span the site's occupation from the Early Preclassic through Late Classic periods[8–12]. Initial dating work at Tikal aimed to articulate the ancient Maya and modern Gregorian calendar systems in order to date Classic period historical events described in glyphic texts on stone monuments across the lowlands[13–15]. To this end, efforts have been directed towards dating wooden lintels and beams from temples throughout the Tikal site core that bear carved dates fixed in the Long Count calendar system. In multiple studies, series of dates were produced from the same lintel in order to document when the wood was cut, carved, and dedicated (e.g.,[13]). Many samples were taken from separate beams to reduce the issues associated with old wood on parts of the lintel.

It should be noted that dates for samples P-235 through P-251 in MesoRAD were originally produced using the Libby half-life of 5568 years rather than the 5730-year half-life conventionally used today. We report corrected dates using the 5730-year half-life and with revised uncertainties following Kennett and colleagues (2013). A total of 56 charcoal samples have also been analyzed from stratified contexts within the Tikal site core at the North Acropolis and Great Plaza. No conventional 14C yr was reported for 21 of these dates[16], p. 989, so they are not considered in the present study. The remaining 35 charcoal dates are all acceptable according to the chronometric hygiene standards, though six dates (P-566, P-563, P-567, P-572, P-573, and P-575) are reported as unacceptable based on their context by[12] and so are not included in our analyses.

#### 3 Supplementary Note 3: Choice of number of mixtures for Maya analysis

We use $K = 10$ finite mixtures for the analysis of Maya radiocarbon dates in the main text. In this section, we briefly validate this choice. Figure 3 shows summaries of the end-to-end Bayesian analysis for all lowland radiocarbon dates (left/red) and just Tikal (right/blue) for $K = 2$, 4, 6, 8, and 10 numbers of finite mixtures. With one important exception, the choice of the number of mixtures does not strongly influence the overall reconstruction. The exception is that without at least $K = 4$ mixtures, the reconstruction for all lowland sites underfits the data. More specifically, it cannot accommodate the Post-Classic resurgence.

Figure 4 demonstrates that even accounting for the influence of sampling and the radiocarbon calibration curve there is support for a Post-Classic resurgence. The upper curve shows the radiocarbon calibration curve fraction modern as a function of calendar date, with the grey bands showing invertible spans and the white bands non-invertible spans. The lower plot shows the end-to-end reconstruction for $K = 2$ (red) and $K = 10$ (green). The histogram in the lower plot is generated from individually calibrating the underlying radiocarbon measurements used in the end-to-end Bayesian inference for All Sites. In particular, for each observation the posterior density over calendar dates is calculated assuming a uniform prior on the calendar date, and for each posterior density the median calendar data is calculated. These median dates are used to generate the histogram.

The descending vertical bars in the lower plot further support the validity of a Post-Classic resurgence, and the need therefore to use at least $K = 4$ mixture components to adequately reconstruct the generating distribution. The descending vertical bars extend two non-invertible regions of the radiocarbon calibration curve. The first has a span of 114.8 years and the second 101.9 years. There are 8 observations with fraction modern values in the first span and 18 with fraction modern value in the second span: 0.070 and 0.18 observations per year, respectively. The median error of observations in the first span is 45 radiocarbon years and in the second span 40 radiocarbon years. Hence, the difference in observations per year cannot be explained by measurement error. Combined with the underlying histogram, this shows that there is indeed a legitimate lull between the Classic peak and the Post-Classic resurgence. At least $K = 4$ mixtures are needed to fit this resurgence, and we chose to use $K = 10$ mixtures for both All Sites and Tikal since it yields a very similar reconstruction to $K = 4$ and allows some additional flexibility reconstructing the generating distribution.

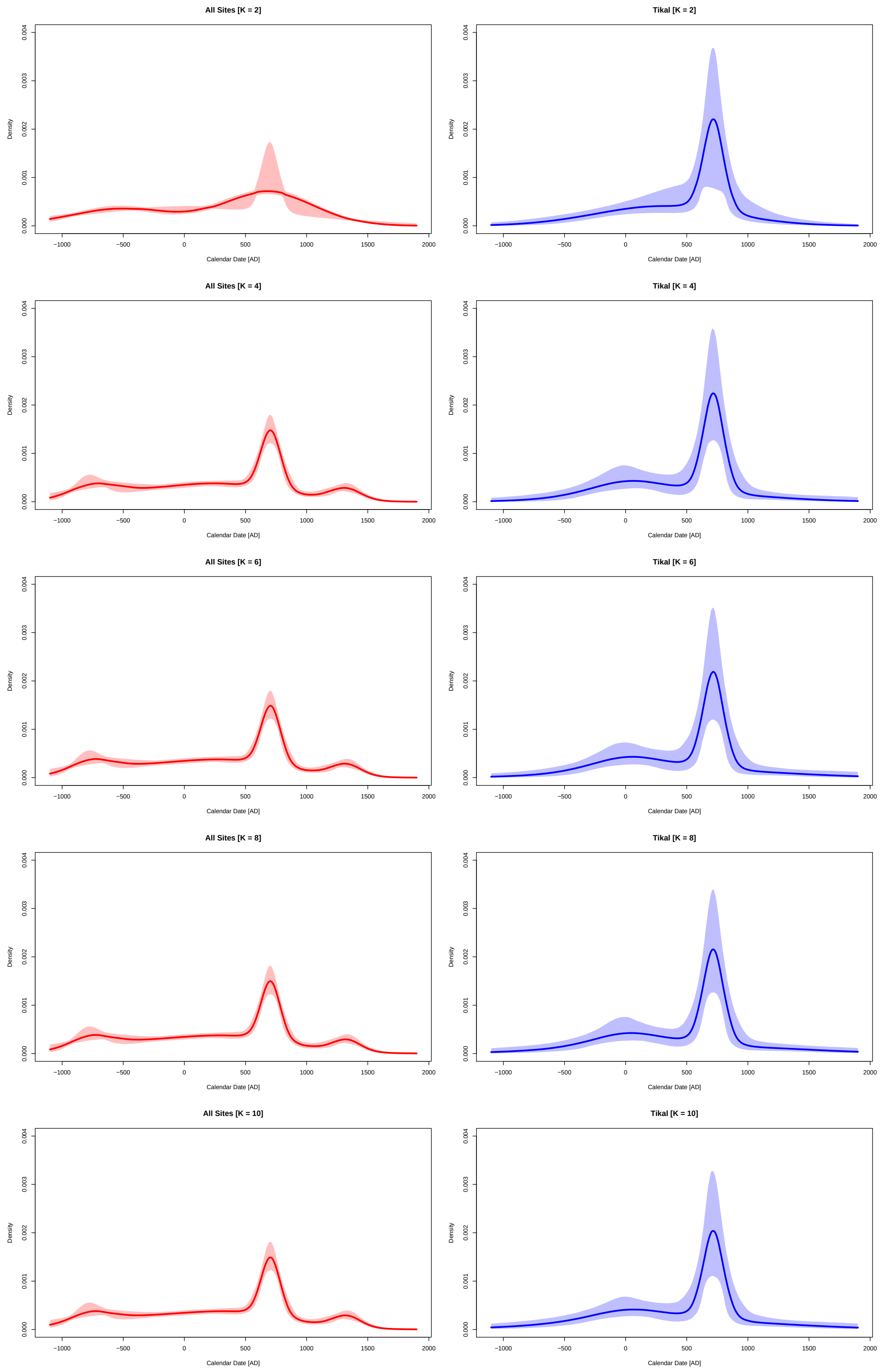

**Supplementary Figure 3.** Summary of end-to-end Bayesian analysis for all dates and just Tikal for $K = 2, 4, 6, 8$, and 10 numbers of mixtures. As in the main text, calendar date is on the x-axis and density on the y-axis, with the solid lines showing the 50% quantiles and the shading the 2.5% to 97.5% quantile bands. The red curves (left column) are for all lowland radiocarbon dates and the blue curves (right column) are for Tikal only.

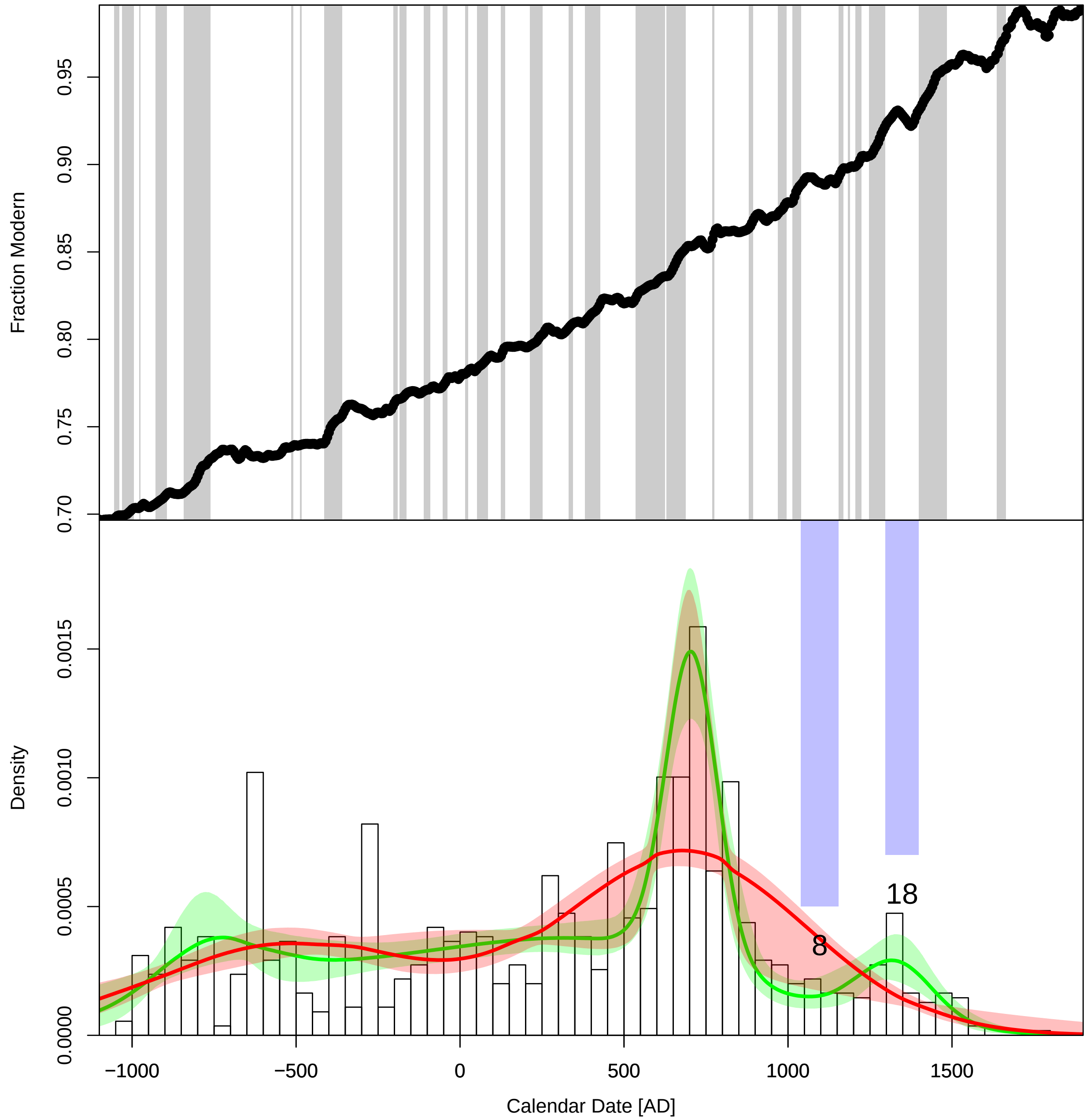

**Supplementary Figure 4.** A more detailed assessment of the end-to-end Bayesian analysis for all dates and $K = 6$ and $K = 10$ numbers of mixtures. The common x-axis is calendar date. The upper plot shows the radiocarbon calibration curve fraction modern on the y-axis and the lower plot probability density. Grey bands in the upper plot delineate spans with a unique, one-to-one mapping between the fraction modern and the radiocarbon date. The histogram in the lower plots is generated directly from individual radiocarbon calibrations of the radiocarbon determinations used in the analysis (in particular, the median calendar date assuming a separate uniform prior for each observation). The red and green curves show the end-to-end reconstructions for, respectively, $K = 6$ and $K = 10$ with the same shading conventions described in Figure 3. The descending vertical bars in the lower plot extend the white, non-invertible spans in the upper plot. While these spans are non-invertible, one can assign observations to these spans (up to measurement and calibration curve uncertainty). The counts for each band are 8 and 18 observations, respectively. The first span is 114.8 years and the second 101.9 years, yielding 0.070 and 0.18 observations per year.
